# Activity-based proteomics uncovers suppressed hydrolases and a *neo*-functionalised antibacterial enzyme at the plant-pathogen interface

**DOI:** 10.1101/2022.12.12.520059

**Authors:** Daniela J. Sueldo, Alice Godson, Farnusch Kaschani, Daniel Krahn, Till Kessenbrock, Pierre Buscaill, Christopher J. Schofield, Markus Kaiser, Renier A. L. van der Hoorn

## Abstract

The extracellular space of plant tissues contains hundreds of hydrolases that might harm colonizing microbes. Successful pathogens may suppress these hydrolases to enable disease. Here, we report the dynamics of extracellular hydrolases in leaves upon infection with *Pseudomonas syringae*.

Using activity-based proteomics with a cocktail of biotinylated probes we simultaneously monitored 171 active hydrolases, including 109 serine hydrolases (SHs), 49 glycosidases (GHs) and 13 cysteine proteases (CPs). The activity of 82 of these hydrolases (mostly SHs) increases during infection, whilst the activity of 60 hydrolases (mostly GHs and CPs) is suppressed during infection. Active β-galactosidase-1 (BGAL1) is amongst the suppressed hydrolases, consistent with production of the BGAL1 inhibitor by *P. syringae*. One of the other suppressed hydrolases, the pathogenesis-related *Nb*PR3, decreases bacterial growth when transiently overexpressed. This is dependent on its active site, revealing a role for *Nb*PR3 activity in antibacterial immunity. Despite being annotated as a chitinase, *Nb*PR3 does not possess chitinase activity, and contains a E112Q active site substitution that is essential for antibacterial activity and is conserved only in *Nicotiana* species.

This study introduces a powerful approach to reveal novel components of extracellular immunity, exemplified by the discovery of the suppression of neo-functionalised *Nicotiana-specific* antibacterial *Nb*PR3.

## INTRODUCTION

Plant pathogens encounter a highly hydrolytic environment when they colonise the extracellular space (apoplast) in plant tissues. Plants secrete hundreds of hydrolytic enzymes (including proteases, glucosidases and lipases), many of which accumulate to high levels during defence. These pathogenesis-related (PR) proteins include chitinases, glucanases, and proteases (Van Loon *et al*., 2006; Doehlemann & Hemetsberger, 2013; Wang *et al*., 2020). In response, most pathogens secrete hydrolase inhibitors when colonising the apoplast to manipulate the host and cause disease. For instance, the oomycete plant pathogen *Phytophthora infestans* secretes Epi1 and Epi10, which are Kazal-like protease inhibitors that suppress defence-related subtilisin-like protease P69B (Tian *et al*., 2004; Tian *et al*., 2005; Tian & Kamoun, 2005). *P. infestans* also secretes cystatin-like protease inhibitors (EpiCs) that target papain-like cysteine proteases (PLCPs) in the tomato apoplast. Tomato PLCPs are also suppressed by Cip1 and Avr2, secreted by the bacterium *Pseudomonas syringae* and the fungus *Cladosporium fulvum*, respectively (Rooney *et al*., 2005; Shabab *et al*., 2008; van Esse *et al*., 2008; Shindo *et al*., 2016). Hydrolase inhibition is not limited to proteases. Soybean pathogen *P. sojae* produces glucanase inhibitor GIP1 (Glucanase Inhibitory Protein 1) (Rose *et al*., 2002) and the fungal corn pathogen *Ustilago maydis* produces PEP1 (Protein Essential during Penetration 1) to inhibit secreted maize peroxidase POX12, thereby preventing the oxidative burst associated with defence (Hemetsberger *et al*., 2012). Taken together, hydrolase inhibition in the apoplast by pathogen-derived molecules is a common infection strategy used to manipulate the host.

We hypothesised that suppressed hydrolases play an important role in immunity. To test this, we used activity-based protein profiling (ABPP), which allows monitoring of protein activities without previous knowledge of substrates or enzyme purification (Cravatt *et al*., 2008; Morimoto & van der Hoorn, 2016; Benns *et al*., 2021). ABPP involves the incubation of a proteome with a chemical probe, which contains a warhead that covalently reacts with the active site; a linker; and a tag to facilitate detection (Morimoto & van der Hoorn, 2016). In combination with analysis by mass spectrometry (MS), ABPP-MS has been applied to study dynamic changes in the activities of serine hydrolases (SHs) in tomato upon infection with *C. fulvum* and *Ralstonia solanacearum*, as well as in *Arabidopsis thaliana* (Arabidopsis) upon infection with *Botrytis cinerea* (Kaschani *et al*., 2009; Sueldo *et al*., 2014; Planas-Marques *et al*., 2017). Here, we greatly increased the power of ABPP-MS with a cocktail of biotinylated probes and taking advantage of a unique model pathosystem.

The interaction between *Nicotiana benthamiana* and the bacterial pathogen *Pseudomonas syringae* pv. *tomato* DC3000 (*Pto*DC3000) provides an ideal system to study hydrolase suppression in the apoplast. The extraction of apoplastic fluid from *N. benthamiana* is a relatively simple procedure, ensuring high yield and purity. *N. benthamiana* is susceptible to the model plant pathogen *PtoDC3000* when it lacks the type-III effector hopQ1-1 (*ΔhQ*), which would otherwise trigger immunity via the Roq1 immune receptor of *N. benthamiana* (Wei *et al*., 2007; Schultink *et al*., 2007). Using the *N. benthamiana*-*Pto*DC3000 model pathosystem and a fluorescent probe labelling retaining ß-glycosidases, we previously discovered the suppression of BGAL1, an apoplast-localised ß-galactosidase (Buscaill *et al*., 2019). BGAL1 participates in flagellin de-glycosylation and thereby plays a major role in the release of the flagellin elicitor that is universally recognised in plants (Buscaill *et al*., 2019). The activity of BGAL1 is suppressed by a small molecule inhibitor produced by *Pto*DC3000 to promote disease (Buscaill *et al*., 2019). This discovery illustrates that suppressed hydrolases play important roles in plant immunity. Here, we explored hydrolase dynamics in apoplastic fluid of infected plants using ABPP-MS with a cocktail of biotinylated probes to uncover additional suppressed hydrolases during infection.

## RESULTS

### The apoplast is rich in hydrolases

To investigate the accumulation of hydrolases in the apoplast of infected and non-infected plants, *N. benthamiana* was infiltrated with water (mock) and *Pto*DC3000(*ΔhQ*). We extracted apoplastic fluids (AFs) at 2 days post infection (2dpi), then digested the proteomes with trypsin and analysed them by LC-MS/MS. We robustly identified 217 proteins carrying a predicted signal peptide (SP, by SignalP, Almagro Armentero *et al*., 2019). Classification of these 217 proteins using PFAM (El-Gebal *et al*., 2019) revealed that 48% of these apoplastic proteins are hydrolases (105 proteins, **Figure 1A**, Supplemental **Table S1**). The other 112 apoplastic proteins are diverse and include eight inhibitors and 13 peroxidases.

**Figure 1.**
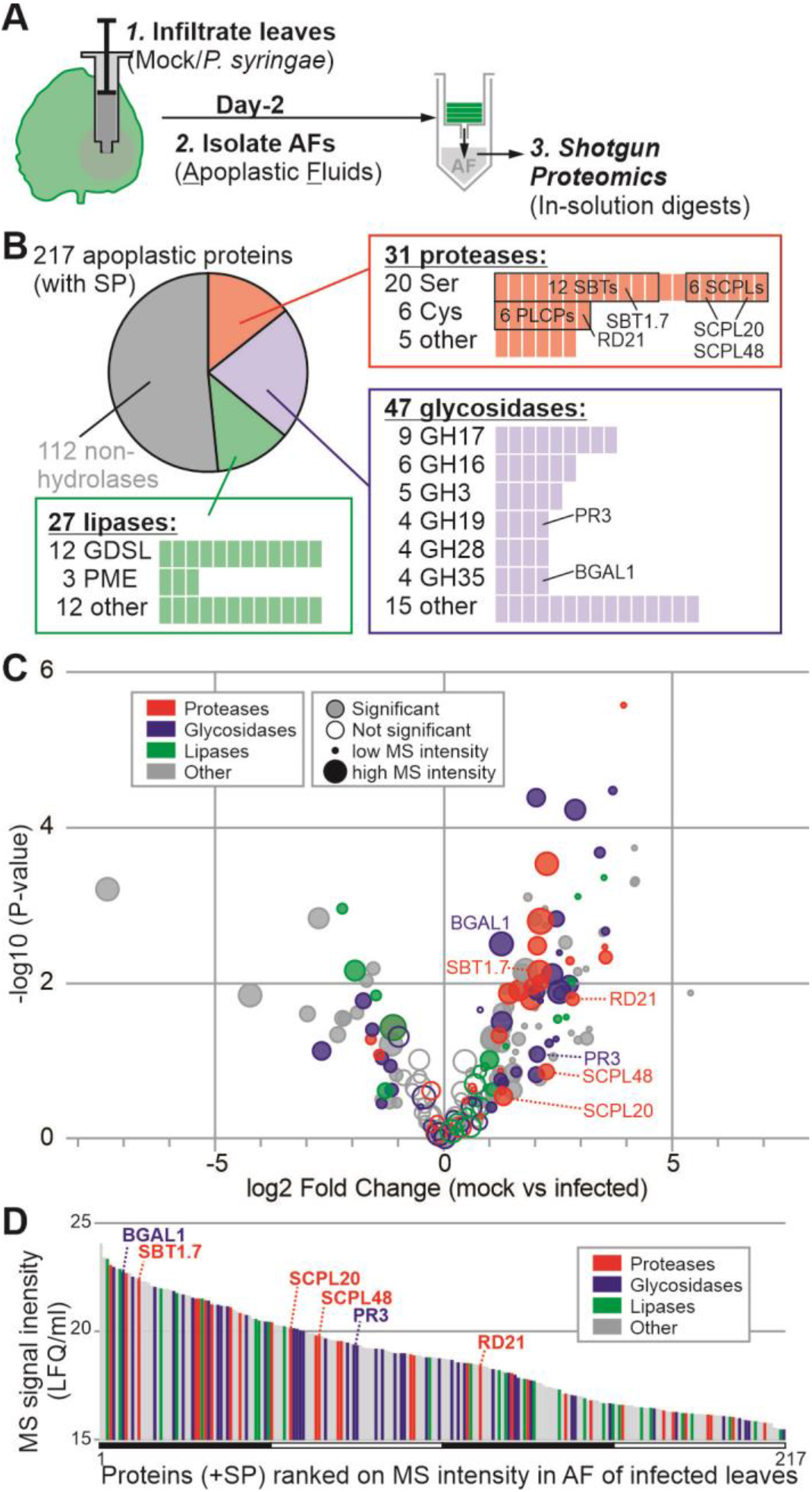
Many plant hydrolases accumulate in the apoplast following infection **(A)** Experimental design. Leaves were inoculated with *Pto*DC3000(*ΔhQ*) (infected) or water (Mock) and apoplastic fluid (AF) was collected at 2 days-post inoculation (2dpi). Proteins were digested with trypsin and analysed by mass spectrometry. **(B)** Many detected secreted proteins are hydrolases. Detected proteins that have a signal peptide predicted by SignalP were annotated with PFAM and classified into proteases (red), glycosidases (blue) and lipases (green), and subdivided into protein families. **(C)** More hydrolytic enzymes accumulate in apoplast upon infection. The p-values of the t-test were plotted against the fold change. The circle diameter reflects the sum of the MS intensities, and filled circles identify proteins with a significant differential accumulation. **(D)** Hydrolytic enzymes are relatively abundant in the apoplast. All 217 SP-containing proteins detected by MS were ranked on average protein intensities detected in AF from mock and infected samples. The different hydrolase classes highlighted in colours and six hydrolases are highlighted. **(A-D)** AF isolated from mock- and *Pto*DC3000(*ΔhQ*)-infected plants were isolated from four independent experiments and analysed by mass spectrometry. LFQ intensities were corrected for protein concentrations measured in the respective AF samples to calculate LFQ/mL. Plant proteins having a predicted SP and detected in all eight samples were retained and used for graphs B-D. See Supplemental **Table S1** for all detected proteins from this ACE_0276 experiment.

The 105 detected hydrolases consist of 31 proteases, 47 glycosidases and 27 lipases (**Figure 1B**). Detected apoplastic proteases belonged to the mechanistic class of Ser proteases (SPs, 20 proteins), Cys proteases (CPs, six proteins), and other proteases (five proteins) (**Figure 1B**). The 20 Ser proteases include 12 subtilisin-like proteases (SBTs, S8 family) and six Ser carboxypeptidase-like proteases (SCPLs, S10). The six Cys proteases are all papain-like Cys proteases (PLCPs, C01). The 47 glycosidases belong to 18 different glycosyl hydrolase (GH) families, which include nine GH17 glycosidases, six GH16 glycosidases, five GH3 glycosidases, and four glycosidases each from families GH19, GH28 and GH35. Finally, the 27 lipases include 12 GDSL-lipases, three pectinacetylesterases (PAEs) and 12 other lipases. This proteome composition is similar to that of a previously reported proteome (Buscaill et al., 2019, ACE_0058).

Protein concentrations in the apoplast can increase 10-fold upon infection (Supplemental **Table S2**), so we corrected our proteomics dataset for this by calculating the MS intensity per mL of apoplastic fluid to facilitate the comparative analysis of protein concentrations. When plotted in a volcano plot, the protein concentrations of abundant hydrolases significantly increase upon infection (**Figure 1D**), consistent with the well-known accumulation of PR proteins, such as chitinases and glucanases (Van Loon et al., 2008). Consequently, apoplastic hydrolases are a major component of the apoplastic proteome of infected plants. When ranked by MS intensity, which is an approximation for protein abundance (Cox *et al*., 2014), 30 hydrolases are amongst the top quartile of most abundant proteins in the apoplast of infected plants (**Figure 1C**). In conclusion, a large proportion of the extracellular proteins encountered by pathogens during infection are hydrolytic enzymes.

### Large-scale activity profiling of the apoplast upon bacterial infection

To investigate changes in hydrolase activities in the apoplast caused by bacteria, we investigated changes in active hydrolases upon bacterial infection with ABPP-MS. To display the active proteome, we labelled the apoplastic proteome with a cocktail of biotinylated activity-based probes including FP-biotin (Liu *et al*., 1999), JJB111 (Chandrasekar *et al*., 2014) and DCG04 (Greenbaum *et al*., 2000) to display the active serine hydrolases (SHs), glycosyl hydrolases (GHs), and cysteine proteases (CPs), respectively. We also included the custom-made Biotin-PD-AOMK (DK-D04, Supplemental **Figure S1A**) to detect vacuolar processing enzymes (VPEs), which have been detected in apoplastic fluids (Sueldo *et al*., 2014). Biotinylated proteins were purified and analysed by MS in n=4 replicates (**Figure 2A**).

**Figure 2.**
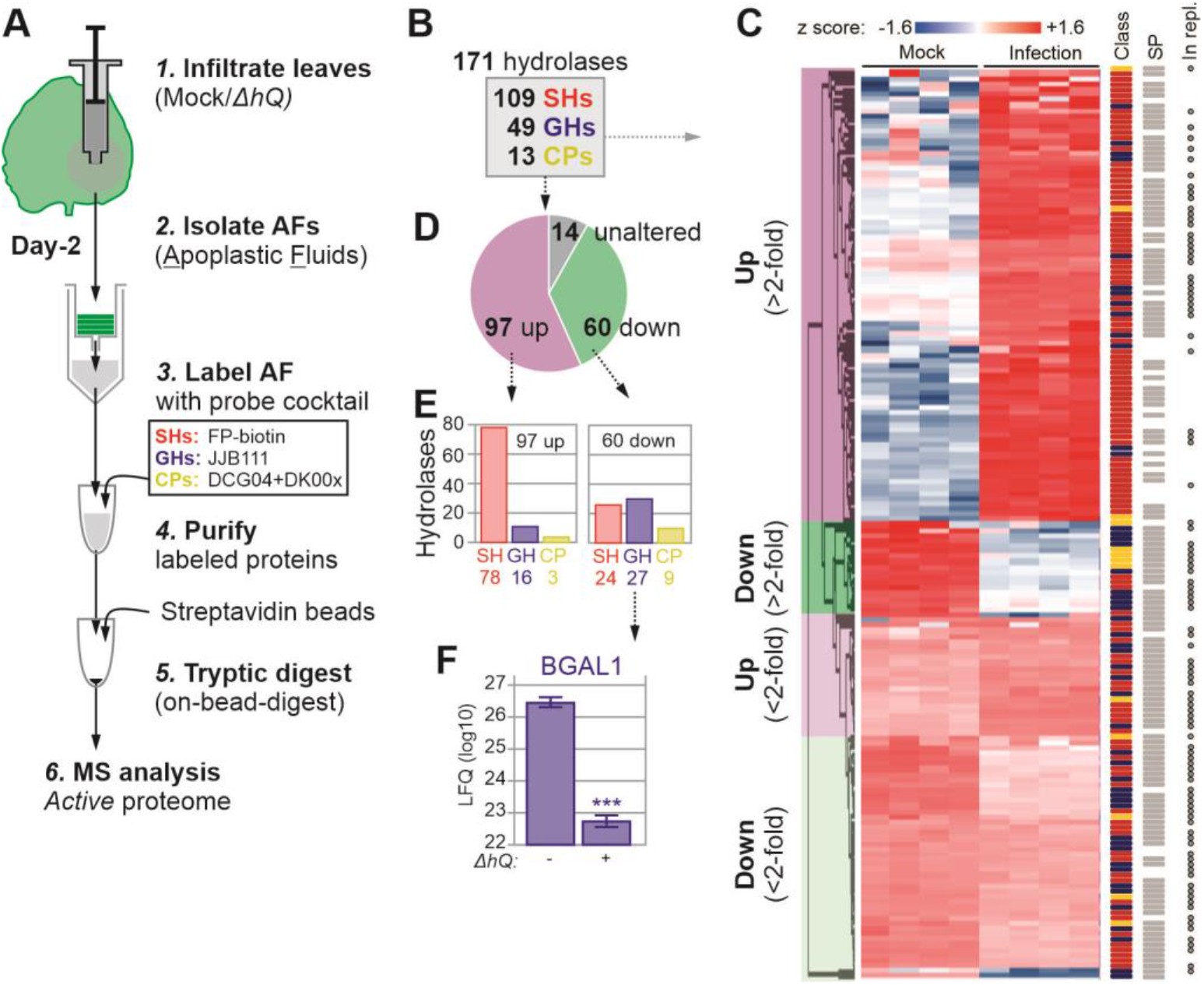
Activity-based proteomics displays hydrolase activity dynamics upon infection. **(A)** Experimental procedure. *N. benthamiana* plants were infiltrated with *Pto*DC3000(*ΔhQ*) or water (Mock) and apoplastic fluid (AF) was collected at 2 days post infection (2dpi). AFs were labelled with a cocktail activity-based probes targeting active serine hydrolases (SHs, red), glycosyl hydrolases (GHs, blue) and cysteine proteases (CPs, yellow). Labelled proteins were purified and identified by MS for n=4 replicates. **(B)** Classification of 171 robustly detected labelled hydrolases. **(C)** Heatmap of the 171 detected active hydrolases detected by activity-based proteomics (experiment ACE136), grouped by category (left) and annotated for being SH/GH/CP; having a SignalP-predicted signal peptide; and for being detected in an independent replicate MS experiment (ACE236) (right). **(D)** Summary of the detected differential active hydrolases showing statistically significant up- or down-regulation upon bacterial infection. **(E)** Summary of the numbers of active hydrolases belonging to each class that are up- or down-regulated. **(F)** BGAL1 labeling is significantly downregulated upon infection. The LFQ intensities for this positive control were extracted from this dataset (ACE_0236).

The annotation of ABPP-MS spectra to the proteome of *N. benthamiana* (Kourelis *et al*., 2019) enabled the detection of 171 predicted probe targets that were all enriched (α=<0.05) compared to the no-probe-control (Supplemental **Table S3**). These probe targets included 109 SHs, 49 GHs and 13 CPs. Overall, 157 target proteins were differentially labelled between mock and *Pto*DC3000(*ΔhQ*) treatments (α=0.05), indicating that 92% of the detected active proteome changes significantly during bacterial infection (**Figure 2C and 2D**). Of the 97 activities that increased upon infection, 78 were SHs, 16 were GHs, and three CPs (**Figure 2E**). Furthermore, we detected 60 reduced hydrolytic activities, including 24 SHs, 27 GHs and nine CPs (**Figure 2E**). Overall, bacterial infection induces mostly active SHs and reduces mostly active GHs and CPs. Importantly, the suppressed hydrolases include BGAL1 (3.67-fold down-regulated, p value = 1.65E-07), as previously described (**Figure 2F**, Buscaill *et al*., 2019). These results indicate that the active apoplastic proteome undergoes large changes during bacterial infection, including 57% of the hydrolases having increased activity, and 35% having reduced activity. Of the 97 increased active hydrolases, 74 had a fold change of at least 2 (FC≥2), consisting of 63 SHs, nine GHs and two CPs. Likewise, 25 of the 60 reduced active hydrolases also had FC≥2, consisting of six SH, 14 GHs, and five CPs.

Although the general trend is that active SHs are induced and active GH and CPs are reduced, we observed many differences within each hydrolase subfamily (**Figure 2E**). We identified 28 active GDSL-like lipases, 23 (most) of which showed increased activity in infected tissues. We also identified 21 subtilisin-like serine proteases (S8, SBTs), of which nine showed decreased activity and nine increased activity upon bacterial infection. Furthermore, we identified 18 serine carboxypeptidase-like proteases (S10, SCPLs) of which twelve had increased activity during infection. We also detected ten carboxylesterases (CXEs) and six pectin acetylesterases (PAEs), all of which were more active upon infection. Of the 17 detected active GH3s, seven were less active in infected tissue. We also detected six active GH79s and six GH35s, including BGAL1 (NbD029635) (Buscaill *et al*., 2019). Eight of the 13 detected papain-like proteases (C01) had a reduced activity upon infection. Both detected VPEs are less active in infected tissue. Altogether, these data demonstrate that the active apoplastic proteome changes drastically upon bacterial infection.

### One of the tested suppressed hydrolases inhibits bacterial growth

An independent experiment confirmed differential activities for 90 of the detected hydrolases (**Figure 3**, Supplemental **Table S4** and Supplemental **Figure S2**). We chose five new hydrolases with a predicted SP (Almagro Armentero *et al*., 2019) that showed robustly suppressed activities upon bacterial infection (**Figure 3**). We selected three SHs (*Nb*SBT1.7, *Nb*SCPL20 and *Nb*SCPL48), one GH (*Nb*PR3) and one PLCP (*Nb*RD21). Because depletion of suppressed hydrolases is less likely to cause disease phenotypes, we tested if increased expression could overcome the suppression and uncover roles of these hydrolases in immunity. We therefore took advantage of the recently developed ‘agromonas’ assay (Buscaill *et al*., 2021), which is based on infections of agroinfiltrated tissues with *Pto*DC3000(*ΔhQ*). We cloned and transiently expressed the five selected hydrolases through agroinfiltration, and infected the agroinfiltrated leaves two days later with *Pto*DC3000(*ΔhQ*). Bacterial growth was determined three days later by plating out dilution series of extracts of infected leaves on selective media. Hydrolase overexpression was confirmed for all the tested hydrolases (Supplemental **Figure S3A**), but only transient expression of *NbPR3* suppresses bacterial growth (**Figure 4A** and Supplemental **Figure S3B**), suggesting a role for *Nb*PR3 in antibacterial immunity. This ‘agromonas’ infection assay has been repeated ten times and *Nb*PR3 increased immunity to *Pto*DC3000(*ΔhQ*) in each of these experiments (Supplemental **Figure S4**). Transient expression of *Nb*PR3 also increased resistance to pv. *tabaci* 6605 (*Pta*6605), but not against pv. *syringae* B728a (*Psy*B728a) (**Figure 4A**), indicating that *Nb*PR3-based immunity might be strain specific.

**Figure 3.**
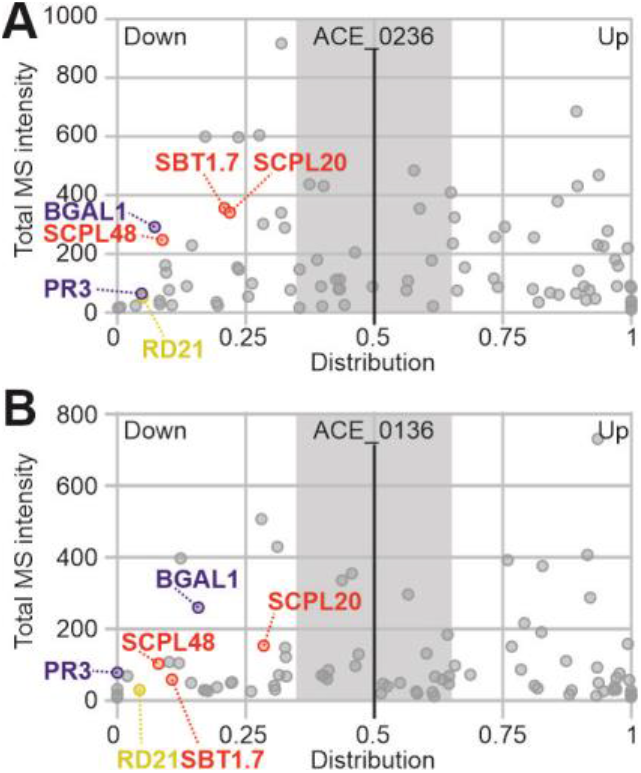
Several secreted hydrolases are consistently suppressed during infection. Distribution graphs of the active proteome from two independent experiments. Data derived from Figure 2 (experiment ACE_0236) **(A)** were compared to a replicate experiment (ACE0136, Supplemental **Figure S3**) **(B)** and used to select consistently suppressed hydrolases. Total MS/MS spectra for mock and infected samples were combined and plotted against the distribution of each protein in mock and infected samples. The distribution was calculated as LFQ_infected_/(LFQ_mock_ + LFQ_infected_). Proteins are represented as dots and only proteins showing the same behaviour in both biological replicates are shown. Six hydrolases (*Nb*SBT1.7a, *Nb*SCPL20, *Nb*SCPL48, *Nb*PR3, *Nb*RD21 and *Nb*BGAL1) are indicated and coloured red, blue and yellow according to hydrolase class (SH, GH and CP, respectively).

**Figure 4.**
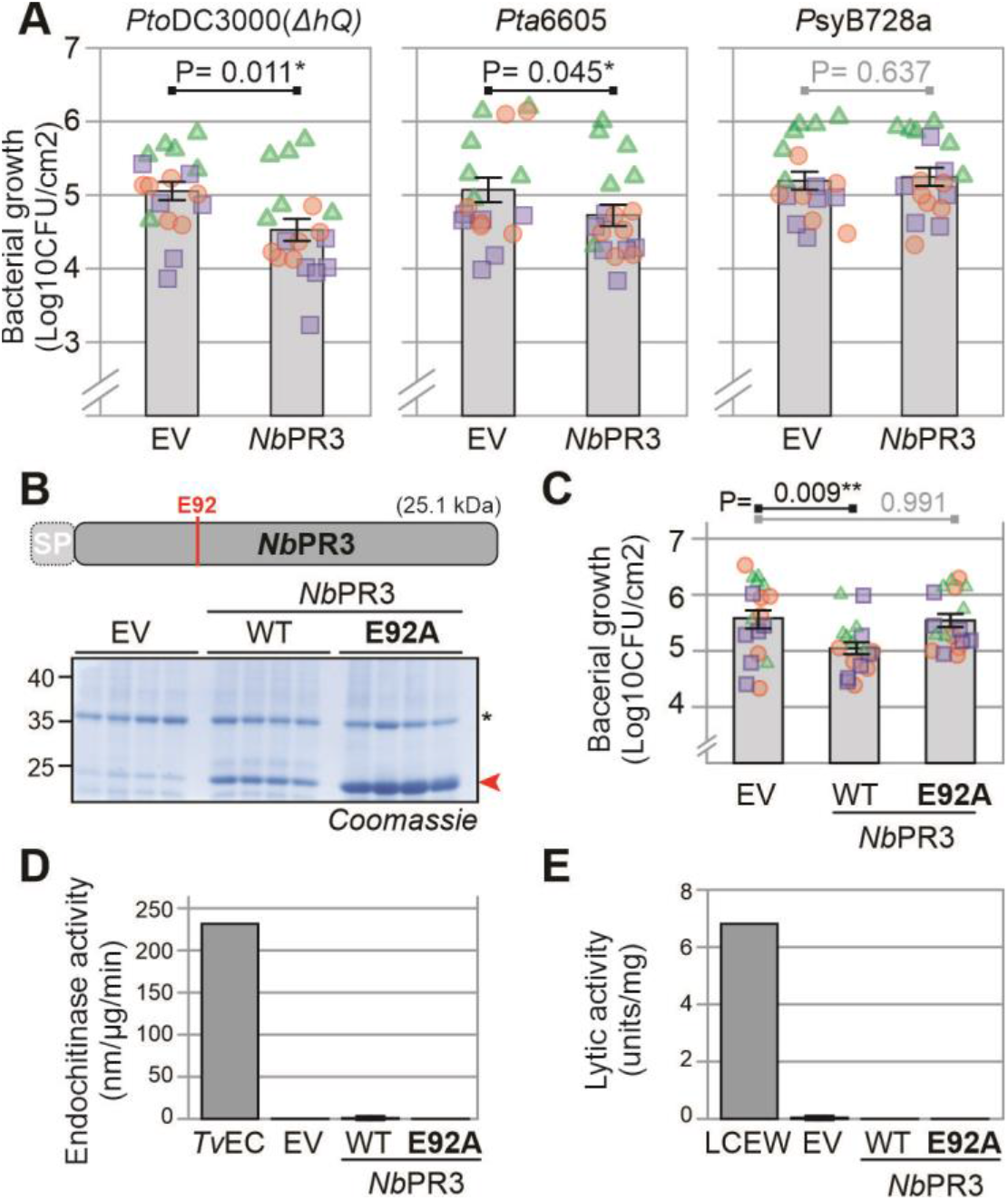
*Nb*PR3 requires its active site to reduce bacterial growth but lacks chitinase and lysozyme activity. **(A)** Transient *Nb*PR3 expression decreases susceptibility to *P. syringae* pathovars *tomato* DC3000(ΔhQ) and *tabaci* 6605, but not *syringae* B728a. *Nb*PR3 and the empty vector (EV) control were transiently expressed by agroinfiltration. Two days later the same leaves were infiltrated with 10^6^ CFU/mL bacteria and bacterial population densities in Log10CFU/cm^2^ were determined after three days. Bars show the mean value of 18 biological replicates performed over three separate experiments, and error intervals represent the standard error. P-values were calculated by two-way ANOVA followed by post-hoc comparison using the Dunnett test to examine the effect of *Nb*PR3 overexpression on bacterial growth. **(B)** The *Nb*PR3(E92) mutant accumulates in the apoplast of agroinfiltrated leaves. *Nb*PR3 and its E92A mutant and the empty vector (EV) control were agroinfiltrated and AF was isolated at day-4 from 4 different replicates, separated on protein gel and analysed by Coomassie staining. The red arrowhead indicates *Nb*PR3 protein. *, endogenous PR2 protein, induced by agroinfiltration. **(C)** Transient expression of *Nb*PR3 but not its catalytic mutant suppresses bacterial growth of *Pto*DC3000(*ΔhQ*). *Nb*PR3, its catalytic site E92A mutant, and the empty vector (EV) control were transiently expressed by agroinfiltration. Two days later the same leaf was infiltrated with 10^6^ CFU/ml *Pto*DC3000(*ΔhQ*) and bacterial population densities in Log10CFU/cm^2^ were determined after three days. Bars show the mean value of 18 biological replicates performed over three separate experiments, and error intervals represent the standard error. P-values were calculated by two-way ANOVA followed by post-hoc comparison using the Dunnett test to examine the effect of *Nb*PR3 overexpression on bacterial growth. **(D)** *Nb*PR3 lacks endochitinase activity. AF from plants transiently expressing *Nb*PR3 or its E92A mutant were incubated with 4-MU-GlcNac3, and the rate of hydrolysis was measured at 355ex/450em and calculated per μg *Nb*PR3 protein estimated by Coomassie staining. The endochitinase of *Trichoderma viride* (*Tv*EC) was included as a positive control. **(E)** *Nb*PR3 lacks lysozyme activity. AF from plants transiently expressing *Nb*PR3 or its E92A mutant were incubated with *Micrococcus lysodeikticus* cells. The change in A450 was measured and converted to units per μg *Nb*PR3 protein estimated by Coomassie staining. The lysozyme of chicken egg white (LCEW) was included as a positive control.

To determine if antibacterial immunity is dependent on *Nb*PR3 activity, we generated an E92A substitution variant of *Nb*PR3, replacing the active site Glu by an Ala residue. This *Nb*PR3^E92A^ protein was successfully expressed upon agroinfiltration (**Figure 4B**), yet was unable to suppress bacterial growth (**Figure 4C**), demonstrating that the intact active site is essential for antibacterial immunity.

*Nb*PR3 belongs to the GH19 hydrolase family and is annotated as an acidic endochitinase. We therefore tested the ability of *Nb*PR3 to degrade a *N*-acetylglucosamine polymer in an *in vitro* fluorogenic reaction using apoplastic fluid from agroinfiltrated leaves expressing either *Nb*PR3 or its E92A mutant as negative control. A commercial chitinase from the fungus *Trichoderma viride* could degrade the substrate, but *Nb*PR3 could not (**Figure 4D**), indicating that *Nb*PR3 does not have endochitinase activity.

The cell wall of bacteria contains peptidoglycan, of which the glycan polymer typically consists of alternating residues of β-(1,4) linked N-acetylglucosamine (NAG) and N-acetylmuramic acid (NAM) units. To test if *Nb*PR3 can hydrolyse peptidoglycan, we monitored the change in the absorbance upon lysis of *Micrococcus lysodeikticus*, which is an established assay for peptidoglycan hydrolysis (Lee & Yang, 2002). However, in contrast to lysozyme from chicken egg white, AF containing *Nb*PR3 culd not lyse the bacteria (**Figure 4E**), indicating that *Nb*PR3 does not have lysozyme activity.

### Nicotiana-specific Q112 is required for antibacterial activity

We next compared the protein sequence of *Nb*PR3 with chitinase-A (CHN-A) from *Nicotiana tabacum*, the closest related sequence with documented endochitinase activity (Suarez *et al*., 2001). Eight residues have been identified in CHN-A as being important for endochitinase activity (Garcia-Casado *et al*., 1998; Tang *et al*., 2004; Chaudet *et al*., 2014; Han *et al*., 2015), only two of which are different in *Nb*PR3 (**Figure 5A**, Supplemental **Figure S5**). Notably, whereas the predicted catalytic general acid Glu residue is present in both sequences (*Nb*PR3^E92^ and CHN-A^E145^), the predicted catalytic general base is absent from *Nb*PR3 (*Nb*PR3^Q112^ vs. CHN-A^E167^). Furthermore, Ser residue S198 in CHN-A is a Thr residue in *Nb*PR3 (T128), though this substitution is common in the plant kingdom. These sequence polymorphisms indicate that *Nb*PR3 may have a catalytic activity that is different from CHN-A, but is still likely to bind carbohydrates because the other residues relevant for chitinase activity are conserved. Indeed, structural modeling indicates that *Nb*PR3 may have a similar fold as chitinase-I (Kezuka et al., 2010), but the region carrying Q112 is different from chitinases (**Figure 5B**).

**Figure 5.**
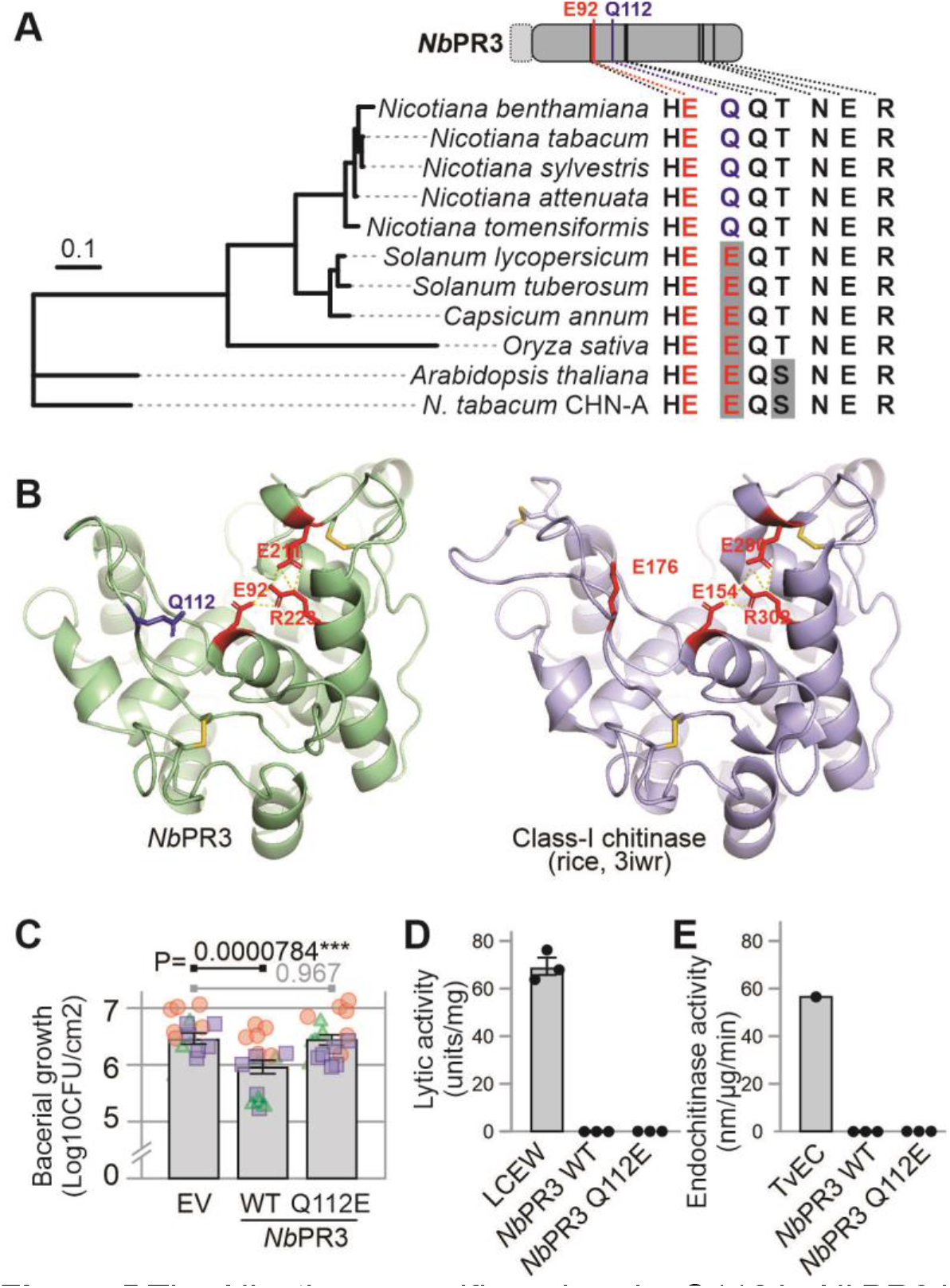
The *Nicotiana-specific* active site Q112 in *NbPR3* is essential for antibacterial activity. **(A)** Conservation of chitinase-relevant residues in *Nb*PR3 homologs. The second catalytic glutamate of chitinases (red) is replaced by glutamine (blue) in all *Nb*PR3 orthologs of *Nicotiana. Nb*PR3 orthologs were identified by BLAST searches, aligned by ClustalO (Supplemental **Figure S6**), and used to construct a maximum likelihood phylogenetic tree. Chitinase A (CHN-A) from *Nicotiana tabacum* was included as the closest related enzyme for which chitinase activity has been demonstrated. The residues relevant for chitinase activity are summarised on the right, showing that the Gln is conserved in *Nicotiana* and is specific for this genus. **(B)** Model of *Nb*PR3 structure, compared to rice class-I chitinase. A model of *Nb*PR3 was generated with SWISS Model, using PDB 3iwr (Kezuka *et al*., 2020) as a template. Catalytically important residues (red), and the noncanonical Q112 (blue), and three cysteine bridges (yellow) are indicated. **(C)** The *Nicotiana*-specific Q112 is essential for antibacterial activity. *Nb*PR3, its Q112E substitution mutant, and the empty vector (EV) control were transiently expressed by agroinfiltration, and two days later the same leaf was infiltrated with 10^6^ CFU/ml *Pto*DC3000(*ΔhQ*). Bacterial population sizes three days later are shown in Log_10_CFU/cm^2^. Bars show the mean value of 18 biological replicates performed over three separate experiments, and error intervals represent the standard error. P-values were calculated by two-way ANOVA followed by post-hoc comparison using the Dunnett test to examine the effect of *Nb*PR3 expression on bacterial growth. **(D)** The Q112E substitution in *Nb*PR3 does not increase lytic activity. AF from plants transiently expressing *Nb*PR3 or its Q112E mutant were incubated with *Micrococcus lysodeikticus* cells. The change in A_450_ was measured and converted to units per μg *Nb*PR3 protein estimated by Coomassie staining. The lysozyme of chicken egg white (LCEW) was included as a positive control. **(E)** The Q112E substitution in *Nb*PR3 does not increase endochitinase. AF from plants transiently expressing *Nb*PR3 or its Q112E mutant were incubated with 4-MU-GlcNac3, and the rate of hydrolysis was measured at 355ex/450em and calculated per μg *Nb*PR3 protein estimated by Coomassie staining. The endochitinase of *Trichoderma viride* (*Tv*EC) was included as a positive control.

Interestingly, Q112 is present in all putative *Nb*PR3 orthologs of the *Nicotiana* genus but is absent in closely related *Solanum* species or Arabidopsis and rice *(Oryza sativa)* (**Figure 5A**, Supplemental **Figure S6**), suggesting that this active site substitution occurred in the *Nicotiana* clade. To test the relevance of Q112 for antibacterial activity of *Nb*PR3, we generated the Q112E mutant and tested its ability to suppress growth of *Pto*DC3000(*ΔhQ*) in the agromonas assay. Importantly, whilst *Nb*PR3 suppresses bacterial growth, the Q112E substitution abolishes this activity (**Figure 5C**), demonstrating that the Q112 residue that is conserved in *Nicotiana* species is relevant for antibacterial activity. The Q112E mutant of *Nb*PR3 does, however, not gain lytic or endochitinase activity (**Figure 5D**), indicating that also other residues contribute to these activities.

### *Nb*PR3 promotes immunity without inducing PR protein accumulation

Previous work on PR-Q, the tobacco ortholog of *Nb*PR3, has revealed that transgenic tobacco overexpressing *PR-Q* also constitutively accumulate PR proteins (Tang et al., 2017). To investigate if *Nb*PR3 expression also induces PR protein accumulation, we monitored the accumulation of PR2, the most abundant secreted PR protein in *N. benthamiana*, upon agroinfiltration with empty vector (EV) and *Nb*PR3 and its E92A and Q122E mutant derivatives. The *NahG*-transgenic *N. benthamiana* (Wulf et al., 2004) was included to determine if PR protein accumulation was dependent on salicylic acid (SA), which does not accumulate in *NahG* transgenic plants. Notably, *Nb*PR3 does not induce PR2 levels and the E92A and Q122E mutants of *Nb*PR3 have similar levels of PR2 when compared to EV and *Nb*PR3 (**Figure 6A**). In fact, expression of *Nb*PR3 and its mutants even reduces PR2 levels (**Figure 6B**), possibly caused by competition on translation and secretion. PR2 levels upon agroinfiltration are much reduced in *NahG*-transgenic plants when compared to WT plants, and again not increased upon *Nb*PR3 expression (**Figure 6**). *Nb*PR3 signals are 2-3 fold higher in the apoplast of plants transiently expressing *Nb*PR3 and derived mutants (**Figure 6B**). Reduced expression of *Nb*PR3 is detected upon agroinfiltration of *NahG* plants, presumably because there is no SA-regulated expression of endogenous *Nb*PR3, and because the 35S promoter used to express *Nb*PR3 contains a SA-responsive *as-1* element (Redman et al., 2002). In conclusion, unlike PR-Q overexpression in tobacco, *Nb*PR3 over expression by agroinfiltration does not induce PR protein accumulation.

**Figure 6.**
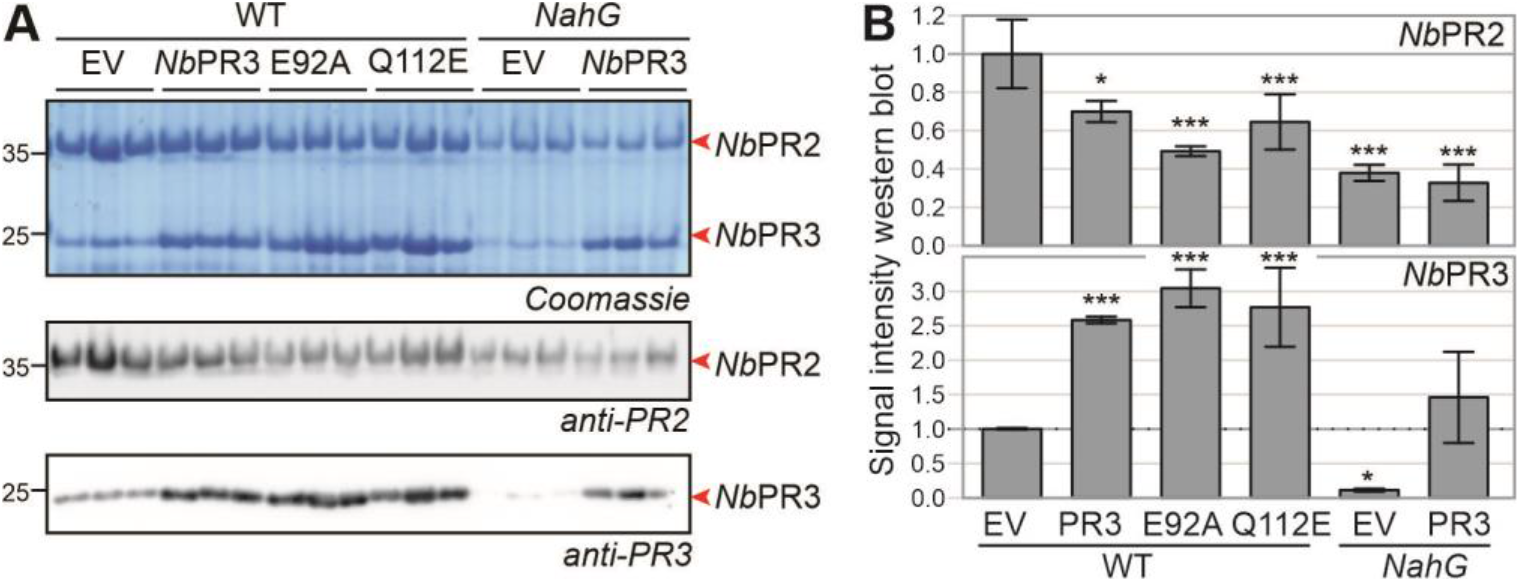
*Nb*PR3 does not promote PR2 protein accumulation. **(A)** *Nb*PR3 and its mutant derivatives were transiently expressed in WT and *NahG*-transgenic *N. benthamiana* in n=3 independent plants. Apoplastic fluids were isolated four days later and analysed by Coomassie staining and western blotting using anti-PR2 and-PR3 antibodies. **(B)** Western blot signals were quantified and normalised for the EV control in WT plants. Error bars represent SD of the n=3 replicates. *, P<0.5; **, P<0.1; ***, P<0.001. P-values are from a one-way ANOVA with Dunnett correction for multiple comparisons.

To investigate if *Nb*PR3 has a direct effect on bacterial growth, we incubated *Pto*DC3000 bacteria in AF isolated from plants expressing EV, *Nb*PR3 and its E92A mutant and monitored bacterial growth over time. Lysozyme mixed with AF of plants agroinfiltrated with EV was included as control. Despite being abundant in AF, *Nb*PR3 did not impact bacterial growth when compared to the EV and E92A controls, whereas lysozyme significantly reduces bacterial growth (Supplemental **Figure S7**), indicating that *Nb*PR3-derived immunity is not direct.

## DISCUSSION

Most SP-containing proteins secreted into the apoplast are hydrolytic enzymes. Here, we explored the apoplast of a model plant-pathogen interaction with activity-based probes and were able to monitor 171 active hydrolases, of which 97 showed increased activity, and 50 had reduced activity upon infection. Disease assays on leaves transiently overexpressing five selected suppressed hydrolases revealed that *Nb*PR3 has antibacterial immunity that requires its active site residue. Despite its annotation as chitinase, *Nb*PR3 has no chitinase or lysozyme activity, presumably because of a E112Q substitution in the active site. This substitution is conserved in *Nicotiana* and is essential for antibacterial activity.

As pathogenic bacteria have adapted to a susceptible host, we anticipate that they will actively suppress secreted hydrolases that may directly or indirectly harm them. Therefore, consistently suppressed hydrolases might be components of the plant immune system that are targeted by pathogen-derived inhibitors. We identified 60 suppressed hydrolases that may play an active role in immunity. Massive changes in protein activities can be caused by altered pH or ion strength, but the influence of these factors was prevented by buffering the AF. Many of the suppressed hydrolases are Cys proteases, which might be suppressed by increased levels of reactive oxygen species (ROS). However, not all Cys proteases are suppressed, indicating that the suppression is caused by selective inhibitors rather than ROS. Indeed, we previously showed that *Pto*DC3000(*ΔhQ*) produced Cip1, which is selectively targeting immune-related Cys proteases of tomato (Shindo *et al*., 2006). Besides Cys proteases, several glycosidases are also suppressed. This includes BGAL1, which is inhibited by a small molecule produced by *Pto*DC3000(*ΔhQ*) (Buscaill et al., 2019). However, given the large proportion of changes, we cannot rule out a general apoplastic regulator (i.e. metabolites, specific ions and post-translational modifications) might influence global protein activities.

Secreted hydrolases that are suppressed during infection can be important novel components of the extracellular immune system of plants. To test this hypothesis, we overexpressed the hydrolases to overcome the threshold of suppression during infection. Indeed, we found that transient overexpression of *Nb*PR3 results in increased immunity to *Pto*DC3000(*ΔhQ*), confirming that supressed hydrolases may act in immunity. Likewise, we found that transient overexpression of BGAL1 increases bacterial resistance (Buscaill *et al*., 2021). However, not all suppressed hydrolases reduced bacterial growth upon overexpression in our agromonas assay. This might be because their overexpression does not overcome the suppression mechanism or because they act in immunity without reducing bacterial growth in the agromonas assay.

The mechanism by which *Nb*PR3 mediates antibacterial immunity, however, remains mysterious. Overexpression of PR-Q in tobacco upregulates multiple defence related genes, an observation which was thought to originate from an unfolded protein response (UPR) caused by the presumed accumulation of unfolded proteins in the ER (Tang et al., 2017). We, however, found that the E92A and Q112E mutants of *Nb*PR3 accumulate equally well but do not trigger immunity, ruling out the UPR being responsible for the observed resistance. That the catalytically important glutamate E92 is essential for the role of *Nb*PR3 in our agromonas assays, implies that catalysis by *Nb*PR3 is required for immunity. However, despite its annotation as an endochitinase, *Nb*PR3 lacks endochitinase activity, at least in our current assays. The absence of bacteriolytic activity indicates that *Nb*PR3 does also not hydrolyse peptidoglycan, unlike Arabidopsis LYS1, a secreted GH18 lysozyme that releases immunogenic fragments from peptidoglycan (Liu *et al*., 2014).

The absence of chitinase and lysozyme activities of *Nb*PR3 is probably caused by the absence of the second catalytic glutamate, which is a glutamine (Q112) in *Nb*PR3. Interestingly, Q112 is a conserved feature of *Nicotiana Nb*PR3 proteins, occurring in all *Nicotiana Nb*PR3 orthologs, including tobacco PR-Q. This indicates that this E112Q substitution must have occurred in a *Nicotiana* ancestor within the solanaceous plant family ~50 million years ago. Importantly, this Q112 substitution is essential for antibacterial activity, indicating that an antifungal PR protein was *neo*-functionalised to gain antibacterial activity. The finding that other residues relevant for chitin binding are still present in *Nb*PR3 indicates that *Nb*PR3 acts on a bacterial glycan.

We discovered that the level of active *Nb*PR3 is consistently downregulated upon infection with virulent *Pto*DC3000(*ΔhQ*) (**Figure 3**), even though the level of the *Nb*PR3 protein increases upon infection (Supplemental **Figure S8**), consistent with being a PR protein. This observation indicates that apoplastic *Nb*PR3 is inhibited during infection, possibly by a small molecule or protein secreted by *Pto*DC3000(*ΔhQ*). The active suppression of an antibacterial enzyme is similar to the suppression of BGAL1 by a small molecule inhibitor produced by *Pto*DC3000(*ΔhQ*) (Buscaill *et al*., 2019), and indicates that exploring the apoplastic battlefield with activity-based proteomics to identify these suppressed hydrolases is an exciting new approach to discover novel components of extracellular immunity in plants.

## MATERIALS & METHODS

### Bacterial infection and apoplastic fluid isolation

*Pto*DC3000(*ΔhQ*) was grown in liquid LB medium (rifampicin 25 μg/mL) overnight 28°C, 220 rpm). The next morning, the bacterial density was measured, and the culture was brought to OD600 = 0.001 with distilled, sterile (MilliQ) water. Cultures were infiltrated in fully expanded leaves of 4-5 weeks-old *N. benthamiana* using a needleless-syringe. Up to two leaves were infected per plant and MilliQ water was used for the mock treatment. Apoplastic fluid was extracted from infected leaves, two days after bacterial infection, as previously described (Joosten, 2012; Hong & van der Hoorn, 2014). Briefly, leaves were harvested and submerged in MilliQ water and ice, with the abaxial side facing down. A vacuum was applied with a pump and subsequently released to allow water intake. Leaves were then rolled into a 50 mL syringe without plunger, placed into a 50 mL tube and centrifuged at 2500 rpm for 25 minutes at 4°C, with slow acceleration and de-acceleration of the rotor. Apoplastic fluid was recovered from the bottom of the 50 mL tube and processed immediately.

### Large-scale labelling and affinity purification

The experiment involved four technical replicates for each biological treatment (mock, infected and no probe control), with each technical replicate corresponding to the apoplastic fluid isolated from 15 plants, two leaves per plant. The experiment was repeated in a subsequent year.

4-5 weeks old *N. benthamiana* plants were infiltrated with *Pto*DC3000(*ΔhQ*) or water (mock) and apoplastic fluid was extracted at 2 dpi as described above. 3 mL of the freshly obtained apoplastic fluid was labelled with the ‘probe cocktail’ (FP-biotin, JJB111, DCG04 and DK-D04) in a reaction mixture containing 50 mM NaAc pH 5, 5 mM DTT and 4 μM of each probe. Labelling was performed at room temperature for 4 hrs with constant rotation. For the no-probe control, an equal volume of DMSO was added to a mixture of 1.5 mL of both AFs. To stop the labelling reaction, proteins were precipitated with chloroform/methanol (Wessel and Flugge, 1984) as follows: 1 volume of ice-cold chloroform, 3 volumes of ice-cold water and 4 volumes of ice-cold methanol were added. The samples were vortexed and subsequently centrifuged (3000 x *g*, 30 minutes, 4°C). The precipitated proteins were resuspended in 2 mL of 1.2% SDS in 1xPBS (Life Technologies) by pipetting, and the solution was further diluted to 0.2% SDS by adding 1xPBS buffer. Proteins were then denatured by heating at 95 °C for 5 minutes. To precipitate labelled proteins, 130 μL of avidin beads (Sigma, A9207) was added to each labelling reaction and incubated for 1 hr at room temperature while rotating, after which the beads were spun down for 10 minutes at 400 x *g*. The supernatant was removed and the beads were washed 5 times with 10 mL of 1% SDS, and then twice with 10 mL of MS-grade water. The beads were then transferred to a protein LoBind tube (Eppendorf, Z666505-100EA).

### Synthesis of DK-D04

All chemicals and solvents were from Sigma-Aldrich, abcr and Fluorochem. Fmoc-Asp-AOMK was synthesised according to a literature procedure (Dolle *et al*., 1994). The synthesis of DK-D04 (I939, Biotin-PD-AOMK, Supplemental **Figure S1A**) was carried out on solid support, according to a general SPPS procedure, following Fmoc-strategy. Couplings have been conducted in a syringe reactor using the corresponding Fmoc-building blocks (3 eq.), HOBt (3 eq.), DIC (3 eq.) and a reaction time of 2 h at room temperature, while the resin suspension was agitated on an orbital shaker. Fmoc-deprotection was achieved by addition of a 5% (v/v) solution of diethylamine in DMF and agitation of the resulting suspension for 15 min. The general washing procedure involved alternated washing of the resin with DMF (3x), MeOH (3x) and DCM (3x). 2-Chlorotrityl resin (1 eq.) was placed in a flame dried flask under an argon atmosphere. Fmoc-Asp-AOMK (4 eq.), dissolved in anhydrous DCM and DIPEA (5 eq.) was added, and the suspension was shaken overnight at room temperature. Methanol was added and stirring was continued for additional 30 min. The resin was transferred into a syringe reactor and washed according to the general procedure. Resin loading was determined by Fmoc-loading test (0.8 mmol/g). For this synthesis, 250 mg (0.2 mmol) of Fmoc-Asp-AOMK loaded 2-Chlorotrityl resin was utilized. A Fmoc-deprotection, and general washing was carried out and a solution of Fmoc-6-Ahx-OH (212 mg, 0.6 mmol), HOBt (81 mg, 0.6 mmol), DIC (76 mg, 0.6 mmol, 93 μL) in DMF (10 mL) was utilized for the next coupling step. Fmoc-deprotection and general washing was carried out and coupling was continued using a solution of Biotin (147 mg, 0.6 mmol), HOBt (81 mg, 0.6 mmol), DIC (76 mg, 0.6 mmol, 93 μL) in DMF (10 mL). The crude product was cleaved off the resin utilizing TFA (3mL) and agitation at room temperature for 1h. The obtained crude product was purified by reversed phase HPLC (H2O:ACN, 0.1% TFA; gradient: from 3 to 80%). The desired product was isolated a colourless powder. Yield: 0.73 mg (0.9 μmol). LC-MS (ESI): m/z = calcd for C_41_H_61_N_6_O_10_S^+^ [M+H]^+^ 829.42, found 829.29. HRMS (ESI): m/z = calcd for C_41_H_61_N_6_O_10_S^+^ [M+H]^+^ 829.41644, found 829.41612.

### Synthesis of TK009

Commercially available E64 Azide (Toronto Research Chemicals, 1 mg, 2.7 μmol, 1 eq.) was dissolved in acetonitrile (100 μL). An aliquot of an aqueous 100 mM CuI solution (16.2 μL, 1.6 μmol, 0.6 eq.) and DIPEA (1.9 μL, 10.8 μmol, 4 eq.) were added. In a second flask, commercially available BDP-630/650-alkyne (Lumiprobe, 5 mg) was dissolved in acetonitrile (100 μL) and an aliquot of this mixture (52.7 μL, 5.4 μmol, 2 eq.) was added to the first reaction solution. The resulting solution was stirred at room temperature for 16 hours after which reaction control by LC-MS indicated completion of the reaction. The desired product was isolated from the solution by injection into a preparative HPLC equipped with a RP-C18 column and run at a flow of 20 mL and with the following gradient program (all solvents contained 0.1% (v/v) TFA): 90% H2O/10% ACN to 30% H2O/70% ACN in 3 minutes, to 25% H2O/75% ACN in 20 minutes. Product-containing fractions were pooled and lyophilized to yield 1.5 mg (64%) of TK009 (I912, Cy5-E64, Supplemental **Figure S1B**). LC-MS (ESI): t_R_ = 10.09 min, m/z calculated for C_42_H_48_BF_2_N_8_O_7_S [M+H]^+^: 857.34, found 857.20.

### On-bead trypsin digestion

The beads were resuspended in 250 μL of 8M urea dissolved in 50 mM Tris-HCl pH8. To reduce disulphide bridges, 12.5 μL of 200 mM DTT was added and beads were incubated at 65 °C for 15 minutes while shaking. The beads were then cooled to 35°C. For the alkylation step, 12.5 μL of 400 mM IAA (iodoacetamide) was added and incubated at 35°C for 30 minutes while shaking and in the dark. Trypsin (Gold Mass spectrometry Grade, Promega) was reconstituted according to manufactures’ instructions and added to the beads. Trypsin digestion was performed overnight at 37°C with shaking. After digestion, the beads were shortly spun down at low speed and the supernatant (containing the peptides) was transferred to a new protein LoBind tube. Beads were washed with 50 μL of MS grade water, spun down and the supernatant was combined with the first supernatant. TFA (trifluoroacetic acid) was added to the peptides to a final concentration of 0.5-1% (v/v). Before mass spectrometry analysis, peptides were purified using Sep-Pak C18 columns (WAT020515, Thermo Fischer) following the instructions provided by the manufacturer.

### Mass spectrometry analysis

Experiments were performed on an Orbitrap Elite instrument (Thermo) coupled to an EASY-nLC 1000 liquid chromatography (LC) system (Thermo) operated in the one-column mode. The analytical column was a fused silica capillary (75 μm × 32 or 36 cm) with an integrated PicoFrit emitter (New Objective) packed in-house with Reprosil-Pur 120 C18-AQ 1.9 μm resin (Dr. Maisch). The analytical column was encased by a column oven (Sonation) and attached to a nanospray flex ion source (Thermo). The column oven temperature was adjusted to 45 °C during data acquisition and at 30 °C in all other modes. The LC was equipped with two mobile phases: solvent A (0.1% (v/v) formic acid, FA, in water) and solvent B (0.1% FA in acetonitrile, ACN). All solvents were of UPLC grade (Sigma). Peptides were directly loaded onto the analytical column with a flow rate around 0.5 – 0.8 μL/min, which did not exceed 980 bar. Peptides were subsequently separated on the analytical column by running a 140 min gradient of solvent A and solvent B (start with 7% (v/v) B; gradient 7% to 35% B for 120 min; gradient 35% to 100% B for 10 min and 100% B for 10 min) at a flow rate of 300 nl/min. The mass spectrometer was set in the positive ion mode and operated using Xcalibur software (version 2.2 SP1.48). Precursor ion scanning was performed in the Orbitrap analyzer (FTMS; Fourier Transform Mass Spectrometry) in the scan range of m/z 300-1500 or 1800 and at a resolution of 60000 with the internal lock mass option turned on (lock mass was 445.120025 m/z, polysiloxane) (Olsen *et al*., 2005). Product ion spectra were recorded in a data-dependent fashion in the ion trap (ITMS) in a variable scan range and at a rapid scan rate. The ionization potential was set to 1.8 kV. Peptides were analysed using a repeating cycle consisting of a full precursor ion scan (1.0 or 3.0 × 10^6^ ions or 30 or 50 ms) followed by 15 product ion scans (1.0 × 10^4^ ions or 50 ms) where peptides are isolated based on their intensity in the full survey scan (threshold of 500 counts) for tandem mass spectrum (MS2) generation that permits peptide sequencing and identification. The collision induced dissociation (CID) energy was set to 35% for the generation of MS2 spectra. During MS2 data acquisition dynamic ion exclusion was set to 120 seconds with a maximum list of excluded ions consisting of 500 members and a repeat count of one. Ion injection time prediction, preview mode for the FTMS, monoisotopic precursor selection and charge state screening were enabled. Only charge states higher than 1 were considered for fragmentation.

### Peptide and protein identification using MaxQuant

RAW spectra were submitted to an Andromeda (Cox *et al*., 2011) search using MaxQuant (version 1.6.10.43) using the default settings label-free quantification and match-between-runs being activated (Cox *et al*., 2014). MS/MS spectra data were searched against the Uniprot *Pseudomonas syringae* pv. *tomato* (strain ATCC BAA-871 / DC3000) (UP000002515_223283.fasta; 5426 entries, downloaded 5/25/2020) reference Proteome and the *Nicotiana benthamiana* database (12864_2019_6058_MOESM10_ESM.fasta; 74802 entries, downloaded 2/21/2020) (Kourelis *et al*., 2019). To estimate the level of contamination, all searches included a contaminants database (as implemented in MaxQuant, 245 sequences) that contains known MS contaminants. Andromeda searches allowed for oxidation of methionine residues (16 Da) and acetylation of the protein N-terminus (42 Da) as dynamic modifications and the static modification of cysteine (57 Da, alkylation with iodoacetamide). The Digestion mode was set to “specific”, Enzyme specificity was set to “Trypsin/P” with 2 missed cleavages allowed, the instrument type in Andromeda searches was set to Orbitrap and the precursor mass tolerance to ±20 ppm (first search) and ±4.5 ppm (main search). The MS/MS match tolerance was set to ±0.5 Da and the peptide spectrum match FDR and the protein FDR to 0.01 (based on target-decoy approach and decoy mode “revert”). Minimum peptide length was 7 amino acids.

The minimum score for unmodified peptides was set to 0. For protein quantification modified peptides (minimum score 40) and unique and razor peptides were allowed. Further analysis and annotation of identified peptides was done in Perseus v1.5.5.3 (Tyanova *et al*., 2016). Only protein groups with at least two identified unique peptides over all runs were considered for further analysis. For quantification we combined related biological replicates to categorical groups and investigated only those proteins that were found in a minimum of one categorical group at least in 3 out of 4 biological replicas. Comparison of protein group quantities (relative quantification) between different MS runs is based solely on the LFQ’s as calculated by MaxQuant (MaxLFQ algorithm). Briefly, label-free protein quantification was switched on, and unique and razor peptides were considered for quantification with a minimum ratio count of 2. Retention times were recalibrated based on the built-in nonlinear time-rescaling algorithm. MS/MS identifications were transferred between LC-MS/MS runs with the “Match between runs” option in which the maximal match time window was set to 0.7 min and the alignment time window set to 20 min. The quantification is based on the “value at maximum” of the extracted ion current. At least two quantitation events were required for a quantifiable protein.

### MS data analysis

Data were analysed with Perseus (Tyanova *et al*., 2016) and only proteins corresponding to *Nicotiana benthamiana* were included in the analysis. Proteins were only considered for analysis if they were identified in three of the four replicates for at least one treatment (i.e., mock or infected). Proteins enriched compared to the no-probe control (NPC) were considered as labelled and furthered analysed. To identify differentially active hydrolases upon infection, we performed a t-test with Benjamini-Hochberg correction for multiple testing (α=0.05) comparing mock and infected samples. Final hydrolase list was manually curated using PFAM (El-Gebal *et al*., 2019) to confirm correct annotation as probe target.

### Cloning of hydrolases

Overexpression constructs for six candidate hydrolases were built by Golden Gate Assembly (Engler *et al*., 2008). The full-length genes were amplified from cDNA using primers summarised in Supplemental **Table S5**. Binary vectors were generated in a Golden Gate reaction with *35S* promoter (pICH51288) and *35S* terminator (pICH41414) into binary backbone pJK001c (Paulus *et al*., 2020) using Bsa1 restriction sites, resulting in binary clones summarized in Supplemental **Table S6**. Binary vectors were transformed into *Agrobacterium tumefaciens* GV3101 (pMP90) by freeze-thawing and selection for kanamycin and gentamicin resistance. The E92A and Q112E mutants of *Nb*PR3 were generated by site-directed mutagenesis using the primers listed in Supplemental **Table S5**, using plasmid pAG001 as a template. The PCR product was digested with DpnI to remove template plasmid and the remaining mutated product was transformed into *E. coli*. Sequence-confirmed positive clones were selected and transformed into *A. tumefaciens*.

### Agroinfiltation

*Agrobacterium* cultures were grown in LB media (kanamycin 50 μg/mL and gentamicin 10 μg/mL) overnight at 28°C. The next day, cultures were spun at 4000 x *g* for 10 minutes at room temperature and resuspended in infiltration buffer (10 mM MgCl2, 10 mM MES, 150 μM acetosyringone) to a final OD_600_0.5. Cultures carrying *Nb*PR3 were co-infiltrated with cultures carrying silencing suppressor P19, and P19 combined with empty vector (EV) was used as negative control. *N. benthamiana* plants were infiltrated at 4-5 weeks-old, and apoplastic fluid was extracted at 4 dpi as indicated above (Joosten, 2012; Hong & van der Hoorn, 2014).

### Protein analysis

Apoplastic fluid was isolated from *N. benthamiana* plants transiently overexpressing hydrolases at 4 dpi. The presence of active serine hydrolases and cysteine proteases was measured by ABPP using the FP-TAMRA probe (Thermo Fisher Scientific) and TK009 (see above), respectively. To label hydrolases, apoplastic fluid samples were incubated in a 50 μl reaction with 0.2 μM FP-TAMRA (for serine hydrolases), or a 250 μl reaction with 2 μM I912 (for cysteine proteases), in the presence of 5 mM DTT and 50 mM sodium acetate pH 5 at room temperature for one hour (serine hydrolases) or four hours (for cysteine proteases). After labelling, the reaction was stopped by precipitation in 4x ice-cold acetone followed by resuspension in 4x gel loading buffer and heating at 90°C for 5 minutes. Samples were separated by SDS-PAGE and visualised using a Typhoon scanner at Cy3 filter (serine hydrolases) or Cy5 filter (cysteine proteases). Coomassie staining was used to check protein loading. *Nb*PR3 was detected by western blot using anti-PR3 antibody designed for tobacco PR3 isoforms (Agrisera; 1:2500 in PBS-T), and an-rabbit-HRP secondary antibody (GE Healthcare).

### Agromonas infection assay

*Agrobacterium* cultures were grown in LB media (kanamycin 50 μg/mL and gentamicin 10 μg/mL) overnight at 28°C. The next day, cultures were spun at 4000 x *g* for 10 minutes at room temperature and resuspended in infiltration buffer (10 mM MgCl2, 10 mM MES, 150 μM acetosyringone) to a final OD_600_ 0.5. All constructs were co-infiltrated with the silencing suppressor P19, and P19 with empty vector was used as mock control. *N. benthamiana* plants were infiltrated when 3-4 weeks-old, and 2-3 fully expanded leaves were infiltrated per plant. Six plants were used per overexpression construct. Plants were covered overnight with a transparent lid to increase humidity. Two days later, agroinfiltrated leaves were infiltrated with 10^6^ CFU/mL *Pto*DC3000(*ΔhQ*) as indicated above. Three days later, leaf discs were punched with a cork borer from each infected leaf, and surface-sterilised with 15% hydrogen peroxide for 2 minutes. Leaf discs were then washed twice in MilliQ and dried. Leaf discs were placed into a 1.5 mL safe-lock Eppendorf tube with three 3 mm diameter metal beads and 1 mL of MilliQ. Tubes were placed in tissue lyser for five minutes at 30 Hertz per second. 200 μL of the lysed tissue was transferred to the first row (A) of a 96-well plate and then serial 10-fold dilutions were made until the last row (20 μL tissue + 180 μL MilliQ water). 20 μL of undiluted tissue and serial dilutions were plated on LB-agar plates containing Pseudomonas CFC Agar Supplement (10 μg/mL cetrimide, 10 μg/mL fucidin and 50 μg/mL cephaloridine, Oxoid SR0103). Plates were allowed to dry, incubated at 28°C for two days and then colonies were counted.

### Endochitinase activity

Apoplastic fluid was used as an enzyme source to test endochitinase activity as described before (Hollis *et al*., 1997; Libantova *et al*., 2009). Briefly, 20 μL of apoplastic fluid was combined with 30 μL of substrate solution (4-methylumbelliferyl-β-D-*N,N′,N″*-triacetylchitotrioside, Sigma) to a final substrate concentration of 18 μM. Reactions were performed in a black 96-well plate and incubated for one hour at 37°C in the plate reader. Measurements were taken every minute for one hour using 355 nm excitation and 450 nm emission. The positive control contained commercial chitinase from *T. viride* at a final concentration of 0.04 mg/mL (Sigma C8241).

### Lysozyme assay

Apoplastic fluid isolated from leaves transiently expressing *Nb*PR3 was used to test lysozyme activity following the instructions of the lysozyme manufacturer. Briefly, 10 μL apoplastic fluid was mixed with 250 μL *Micrococcus lysodeikticus* cells resuspended in potassium phosphate buffer at 0.15 mg/mL. Absorbance at 450 nm was recorded in Tecan plate reader at 15 second intervals for five minutes at 25°C. The maximum linear rate of A_450_ decrease was calculated and converted into enzymatic units per μg protein. The positive control contained 0.1 μg commercial lysozyme from chicken egg white (Sigma L6876).

## Acknowledgements

We thank Ursula Pyzio for excellent plant care; Sarah Rodgers and Caroline O’Brian for technical assistance; and Friederike Grosse-Holz, Jiorgos Kourelis and Mariana Schuster for constructive discussions. We thank Sylvestre Marillonnet and Nicola Patron for providing pICH41414 and pICH51288 via Addgene; Jonathan Jones for providing seeds of *NahG*-transgenic *N. benthamiana*; and Hermen Overkleeft for providing JJB111 until 2019. This project was financially supported by the European Research Council grant (ERC-AdG-2020) 101019324 ‘ExtraImmune’; the BBSRC research grant ‘GH35’ (BB/R017913/1); and the Interdisciplinary Doctoral Training Program (DTC) of the BBSRC (DDT00060).

## Author contributions

SLD, AG and RH planned and designed the research; SLD, AG and PB performed experiments; FK and MK performed proteomic analysis; DK, TK, CJS and MK synthesised probes; SLD and RH wrote the manuscript with help of all authors. SLD and AG contributed equally.

## Data availability

The mass spectrometry proteomics data have been deposited to the ProteomeXchange Consortium via the PRIDE partner repository (Perez-Riverol *et al*., 2022).

## Supplemental Files

**Figure S1.**
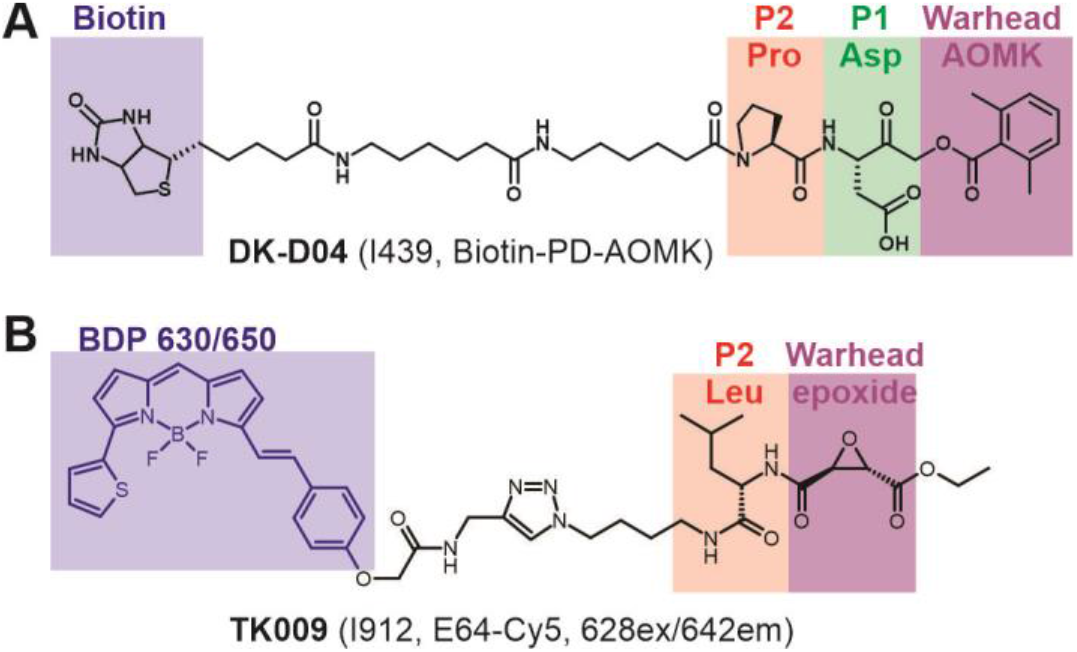
Structure of additional activity-based probes. **(A)** Structure of DK-D04 (Biotin-PD-AOMK), a probe for biotinylation of VPEs. **(B)** Structure of TK009 (Cy5-E64), a probe for fluorescent labelling of PLCPs.

**Figure S2.**
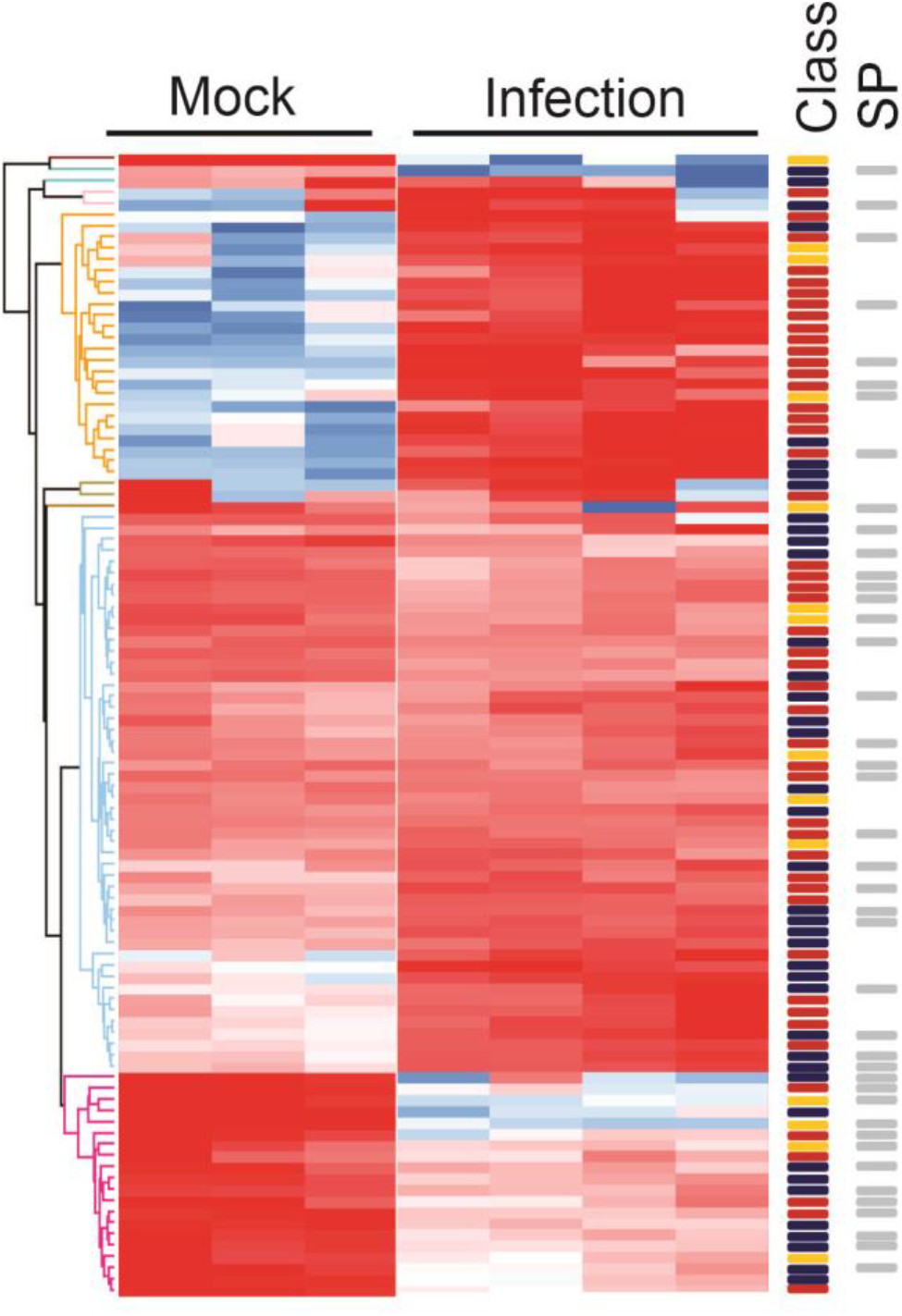
Replicate activity-based proteomics experiment. Heatmap of 105 active hydrolases detected by activity-based proteomics (experiment ACE136), grouped by category (left) and are annotated as SH(red) / GH(blue) / CP(yellow); hydrolases with a SignalP-predicted signal peptide are marked in grey (right). *N. benthamiana* plants were infected with *Pto*DC3000(*ΔhQ*) or water (mock); apoplastic fluid was collected at 2 days post infection (2dpi). Apoplastic fluid was labelled for active serine hydrolases (SHs), glycosyl hydrolases (GHs) and cysteine proteases (CPs) followed by affinity purification of the labelled proteins and MS analysis.

**Figure S3.**
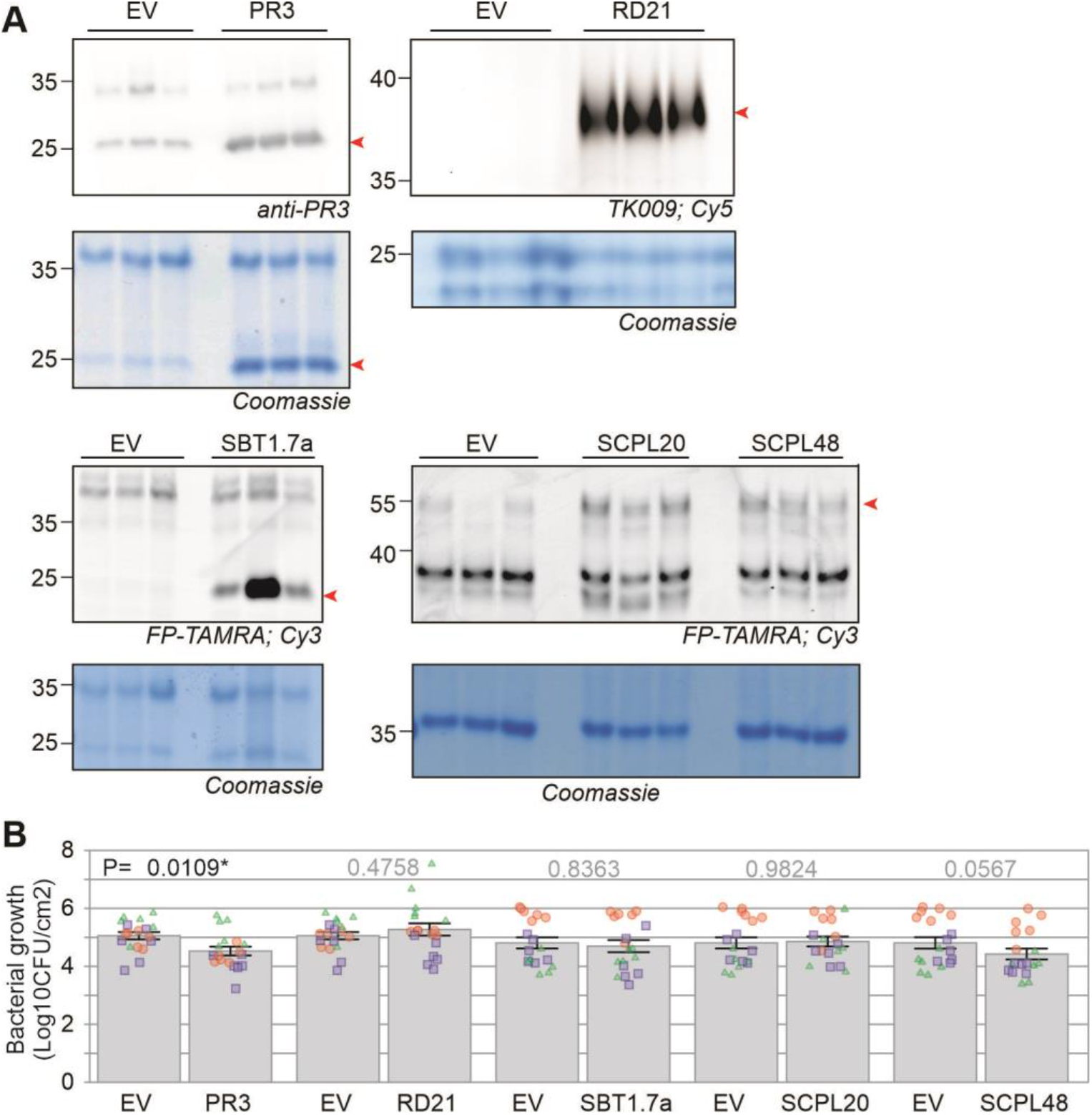
Transient hydrolase expression and agromonas assays. **(A)** Confirmation of hydrolase expression. Hydrolase genes were transiently overexpressed by agroinfiltration of *N. benthamiana*. AF was isolated in triplicate at day-4 and labelled with fluorescent probes selective for papain-like proteases (TK009) or Ser hydrolases (FP-TAMRA). Labelled samples were separated on protein gels, which were scanned for fluorescence and stained with Coomassie Blue. *Nb*PR3 accumulation was detected with an anti-PR3 antibody and detected by Coomassie Blue staining. **(B)** *Nb*PR3 reduces bacterial growth. Hydrolases were transiently overexpressed by agroinfiltrations. At day 2, agroinfiltrated leaves were infiltrated with 10^6^ CFU/ml *Pto*DC3000(*ΔhQ*). At day 5, leaf extracts were generated, diluted and spotted on plates containing CFC antibiotics to select for *P. syringae*. Bacterial growth is shown in Log10CFU/cm^2^. Bars show the mean value of 18 biological replicates performed over three separate experiments, and error intervals represent the standard error. P-values were calculated by two-way ANOVA followed by post-hoc comparison using the Dunnett test to examine the effect of hydrolase overexpression on bacterial growth.

**Figure S4.**
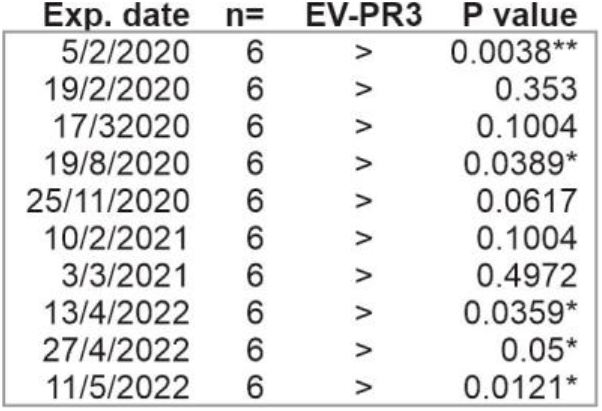
Replicates of infection assays show increased resistance upon *Nb*PR3 expression. *Nb*PR3 and the empty vector (EV) control were transiently expressed by agroinfiltration. Two days later the same leaf was infiltrated with 10^6^ CFU/ml bacteria and bacterial population densities were determined three days later, expressed Log_10_CFU/cm^2^. Summarised are 10 experiments, each having n=6 biological replicates. Bacterial growth on *Nb*PR3 expressing leaves was always lower (>) than the EV control, and statistically significant for 5 of the 10 experiments.

**Figure S5.**
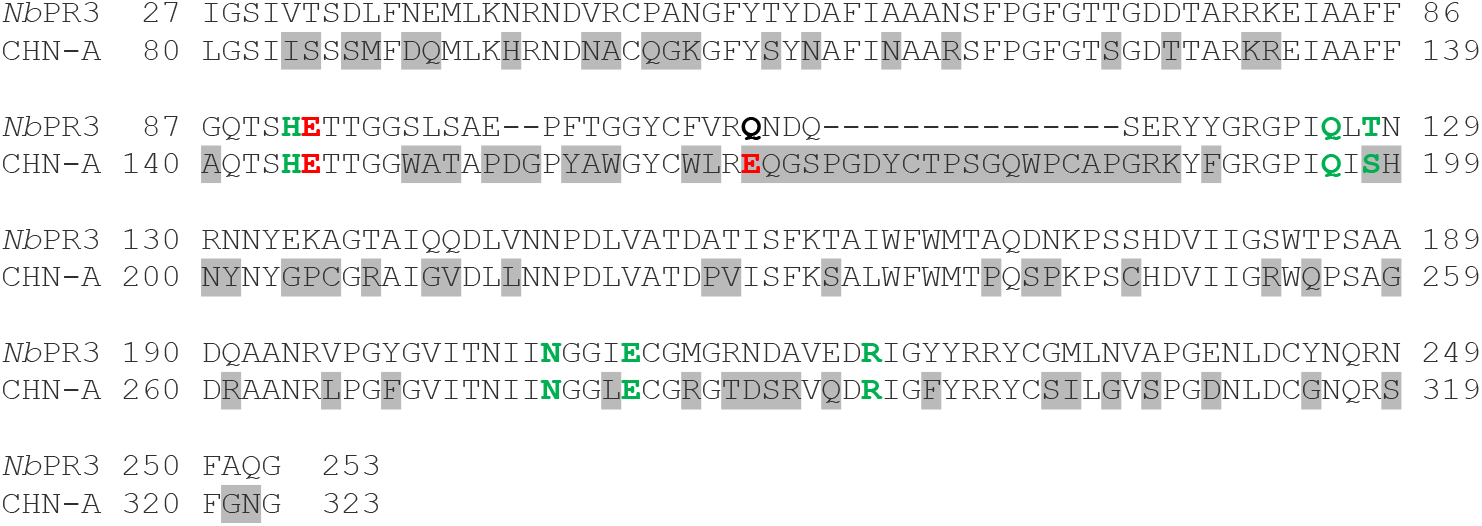
Protein sequence alignment of *Nb*PR3 and endochitinase CHN-A. The mature *Nb*PR3 protein sequence was aligned with the class I endochitinase CHN-A from *N. tabacum*. Residues that differ from *Nb*PR3 (grey), key catalytic residues (red) and other residues relevant for chitinase activity (green) are highlighted.

**Figure S6.**
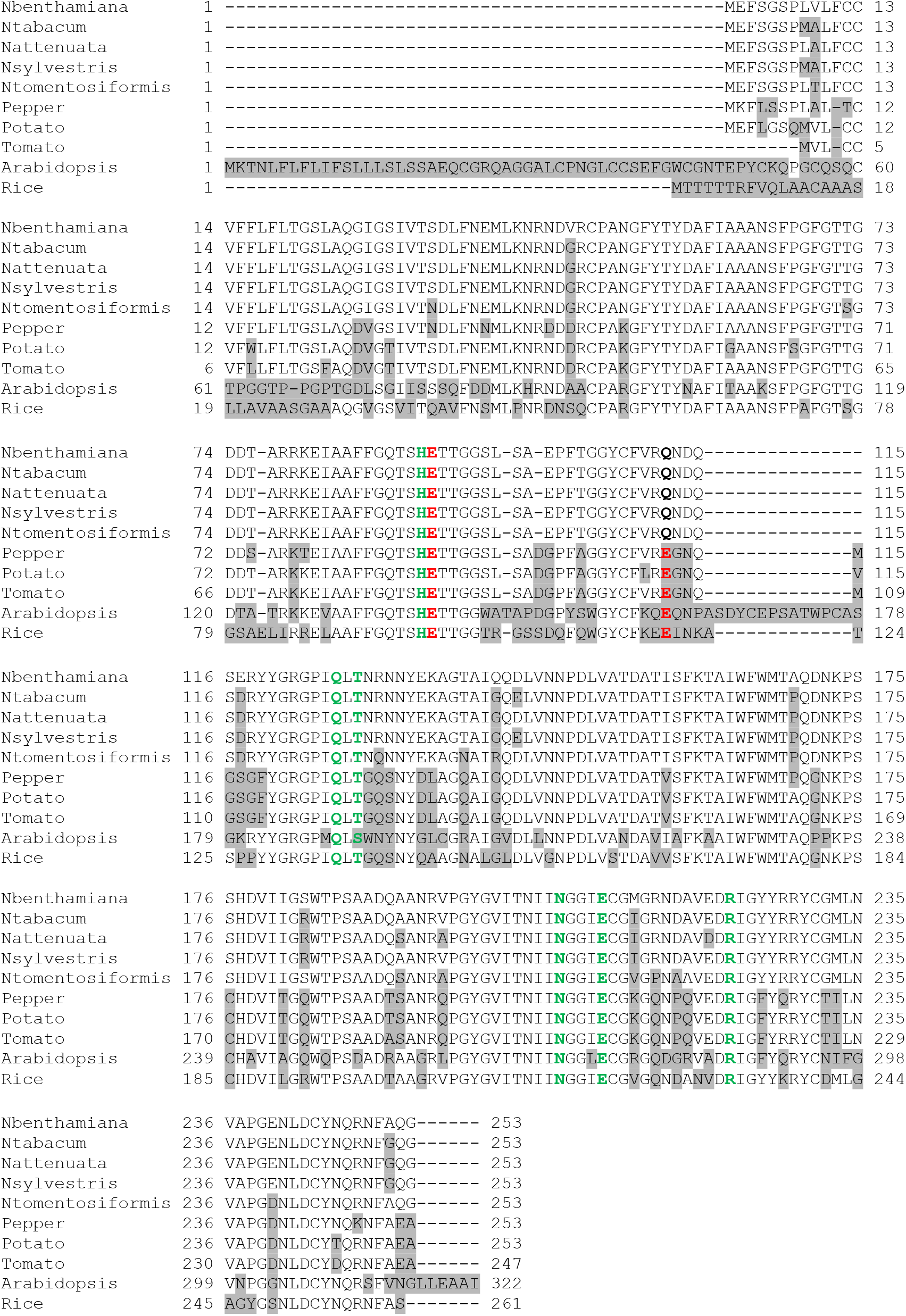
Protein alignment of putative *Nb*PR3 orthologs from diverse plant species. ClustalO alignment of putative *Nb*PR3 orthologs from several *Nicotiana* species, pepper, potato, tomato, *Arabidopsis thaliana* and rice. Differences with *Nb*PR3 (grey), key catalytic residues (red) and other residues relevant for chitinase activity (green) are highlighted.

**Figure S7.**
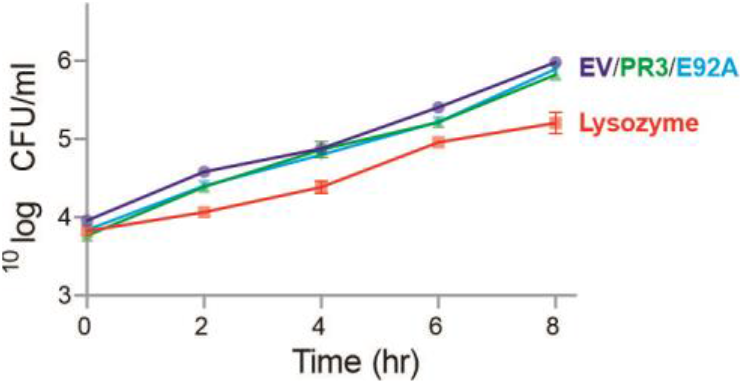
*Nb*PR3 does not impact bacterial growth *in vitro*. AF from agroinfiltrated plants expressing *Nb*PR3 or its E92A mutant or the empty vector (EV) control, with or without added lysozyme, was incubated with bacteria and samples were taken at various time points and colony forming units (CFU) was determined by plating out dilution series.

**Figure S8.**
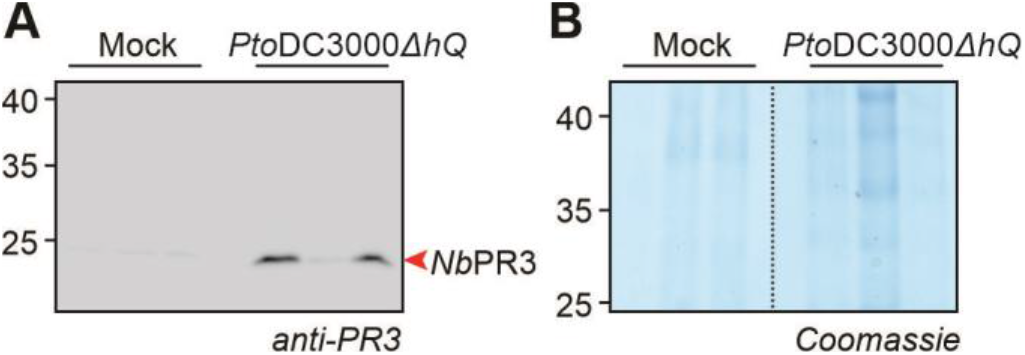
*Nb*PR3 accumulates in the apoplast upon infection. *N. benthamiana* leaves were infiltrated with water (mock) or *Pto*DC3000(*ΔhQ*) and AF was isolated from three biological replicates at 2dpi and equal volumes were analysed by western blot with the anti-PR3 antibody **(A)** and on gel, stained with Coomassie Blue **(B)**.

**Table S1** In-solution digest (ACE_0276)

**Table S2.**
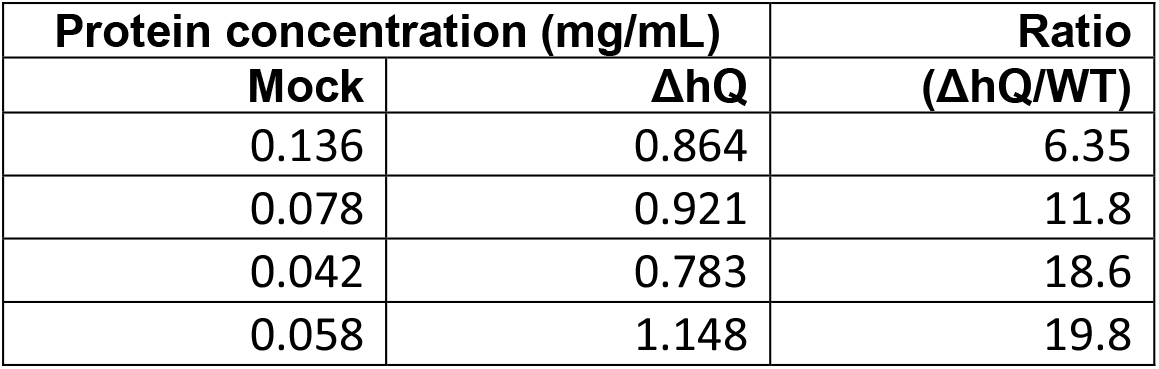
protein concentrations in AF.

| Protein concentration (mg/mL) |  | Ratio |
| --- | --- | --- |
| Mock | ΔhQ | (ΔhQ/WT) |
| 0.136 | 0.864 | 6.35 |
| 0.078 | 0.921 | 11.8 |
| 0.042 | 0.783 | 18.6 |
| 0.058 | 1.148 | 19.8 |

**Table S3** On-bead digest (ACE_0237)

**Table S4** On-bead digest (ACE_0137)

**Table S5.**
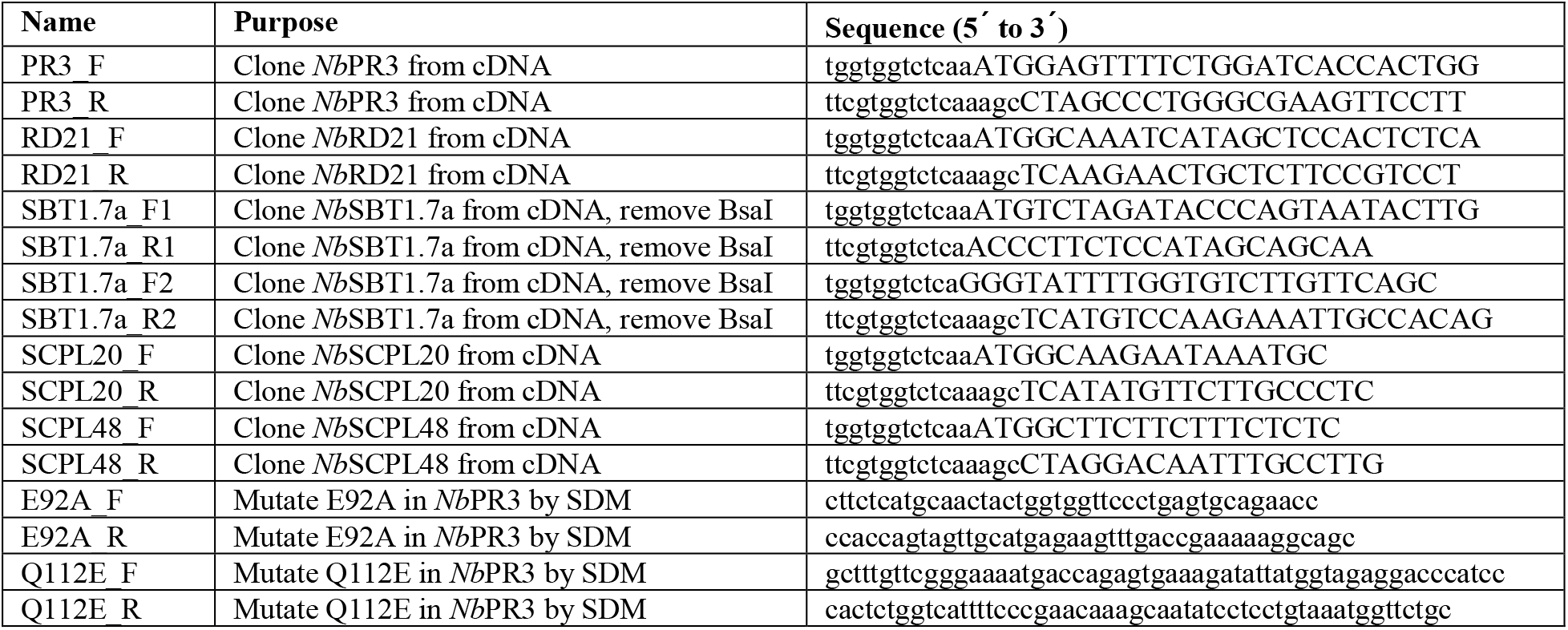
Used oligonucleotides.

**Table S6.** Used plasmids.

| **Name** | **Description** | **Reference** |
| --- | --- | --- |
| pICH51288 | pL0M-PU-35S-TMV-3-5128835S | Engler *et al.*, 2014 |
| pICH41414 | pL0M-T-35S-1-4141435S | Engler *et al.*, 2014 |
| P19 | Binary 35S::P19 | Van der Hoorn *et al.*, 2003 |
| pJK001c | Binary empty vector (EV) | Paulus *et al.*, 2020 |
| pAG001 | Binary 35S::NbPR3 | This work |
| pPB034 | Binary 35S::NbRD21 | This work |
| pAG003 | Binary 35S::NbSBT1.7a | This work |
| pAG004 | Binary 35S::NbSCPL20 | This work |
| pAG005 | Binary 35S::NbSCPL48 | This work |
| pAG006 | Binary 35S::NbPR3<sup>E92A</sup> | This work |
| pAG007 | Binary 35S::NbPR3<sup>Q112E</sup> | This work |

